# Continuity within somatosensory cortical map shapes the integration of optogenetic input

**DOI:** 10.1101/2021.03.26.437211

**Authors:** H. Lassagne, D. Goueytes, D.E Shulz, L. Estebanez, V. Ego-Stengel

## Abstract

The topographic organization of sensory cortices is a prominent feature, but its functional role remains unclear. Particularly, how activity is integrated within a cortical area depending on its topography is unknown. Here, we trained mice expressing channelrhodopsin in cortical excitatory neurons to track a bar photostimulation that rotated smoothly over the primary somatosensory cortex (S1). When photostimulation was aimed at vS1, the area which contains a contiguous representation of the whisker array at the periphery, mice could learn to discriminate angular positions of the bar to obtain a reward. In contrast, they could not learn the task when the photostimulation was aimed at the representation of the trunk and legs in S1, where neighboring zones represent distant peripheral body parts, introducing discontinuities. Mice demonstrated anticipation of reward availability, specifically when cortical topography enabled to predict future sensory activation. These results are particularly helpful for designing efficient cortical sensory neuroprostheses.

**Teaser:** Optogenetic stimulation sweeping the cortical surface: A way to provide precise sensory information and guide behaviour.

## Introduction

Primary sensory areas of the neocortex are involved in sensory perception for several modalities. For example, microstimulations of the cortex of human participants produce vivid sensory percepts, whether visual, tactile or auditory (*1*–*4*).

One striking characteristic that was revealed already by early reports, using microstimulations or recording techniques, has been that these sensory areas are organized topographically with respect to the periphery (*1*, *5*, *6*). In the primary somatosensory cortex (S1), Penfield and collaborators have shown that neighboring cortical zones encode information from neighboring patches of skin on the body (*1*). Such cortical representation of the body surface in S1 is a common feature of mammals. In rodents, the map corresponds to an anatomical organization of the cortical network, in which distinct clusters of cells, called barrels, correspond to the peripheral whiskers on the snout (*7*).

To what extent direct activation of cortical ensembles in these maps may be necessary and sufficient for eliciting sensory percepts is an area of active research (*8*–*10*). Animals can be trained to report direct cortical stimulation of primary sensory areas for all modalities (*9*–*14*). Interestingly, several studies have shown that animals trained in a sensory perception task could rapidly generalize when the peripheral stimuli were replaced by cortical microstimulations (*9*, *14*–*20*). This seemingly immediate interchangeability of natural and artificial stimuli strongly supports a prominent role of cortical activity in sensory perception.

Throughout these experiments, stimulation of distinct cortical zones in a topographic sensory map elicited distinct percepts matching the expected peripheral locations on the sensory organ (*9*, *10*, *21*– *23*). This consistency has been interpreted as additional evidence that the topographical organization of sensory areas serves a fundamental function for sensory processing (*24*). Alternatively, topography could be a mere consequence of the way cortical areas evolved and develop in the organism. In this opposing view, the spatial arrangement of cortical zones as an orderly mosaic may have no functional impact on the computations performed by the cortex (*25*).

To date, most studies linking cortical activity and perception have focused on single-stimulus detection, or discrimination between several stimuli each presented individually. In such conditions, cortical zones may indeed be read out sequentially and independently of the global map in which they are embedded. More complex stimuli in which information is spatially and temporally distributed are necessary to reveal the impact of topography on cortical computations and perception. Indeed, functional topography is known to be intrinsically associated with precise intracortical connectivity patterns (*26*–*28*). These highly non-random connections could result in differential sensory processing depending on the ensembles of neurons being activated.

In line with this view, we hypothesize that the detailed structure and connectivity of the cortex shapes the integration of stimuli in nonlinear ways (*29*) and that this impacts sensory perception. In particular, patterns of cortical activation matching expected activation from spatiotemporal continuous patterns in the periphery could be favored by the cortical circuitry, allowing fast interpretation by the animal and even temporal expectancy (*30*). Beyond a better understanding of sensory perception, uncovering such computational rules could be critical to design the efficient delivery of sensory information directly into neuronal networks, an endeavor actively pursued for sensory neuroprostheses and closed-loop brain-machine interfaces (*9*, *31*–*33*).

In this study, we focus specifically on the functional significance of map topography for sensory processing and perception. We chose to causally manipulate the activity of cortical neurons by patterned optogenetic activation, so that the behavior of the animal was necessarily due to perception of cortical activity. We designed a continuous moving bar stimulus projected onto a cortical surface known for either its orderly two-dimensional topography (the barrel cortex, vS1), or the presence of discontinuities in its topography (the trunk and legs representation, bS1), or its lack of a clear topography (the posterior parietal cortex, PPC). The capacity of the mouse to perceive this cortical stimulus and anticipate its trajectory depended on the cortical area, pointing to differential sensory processing.

## Results

### Mice learn the angular position of a rotating optogenetic stimulus projected onto the barrel cortex

In this study, we tested whether head-fixed, water-restricted mice expressing channelrhodopsin in excitatory neurons could actively track the position of an optogenetic stimulus applied on the representation of whiskers in the primary somatosensory cortex (Fig. 1A-B). We first determined the location of at least three barrels, including the C2 barrel, by intrinsic imaging (Fig. 1C). The locations were confirmed postmortem by histological barrel map reconstruction (Fig. 1D, (*34*)). We then rotated a light bar stimulus inside a disk centered on the representation of whisker C2 (Fig. 1E). For each trial, the light bar turned smoothly according to a predetermined trajectory (see Methods and Fig. 1F-H), thus activating sequentially contiguous zones of the barrel cortex. Mice could lick for water reward when the photostimulation bar was in a Rewarded zone (green area, Fig. 1E). Licking when the bar was in the No lick zone (red area) immediately ended the trial and started a 5 s intertrial interval during which no stimulus was applied. Thus, mice had to learn both to lick for reward and refrain from licking, depending on the angular position of the bar.

**Fig. 1.**
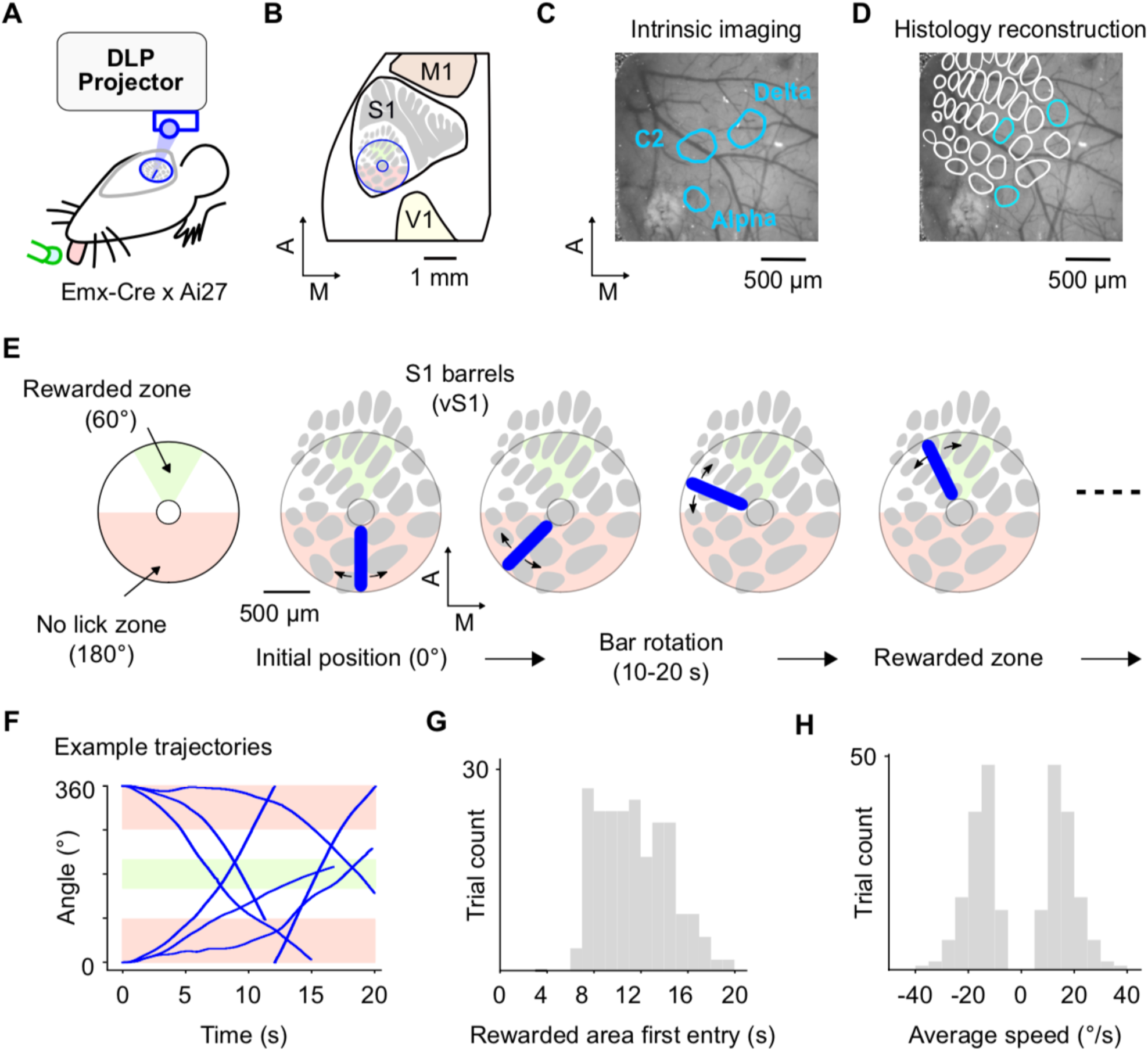
Mice were trained to lick for reward when a moving photostimulation bar entered a defined vS1 zone. **(A)** Sensory-guided licking task. A digital projector sends frames through an optical window, stimulating the mouse cortex. A water tube with a capacitive sensor detects licks and delivers rewards when appropriate. **(B)** Location of the stimulation disk over vS1. The barrel map was adapted from (*54*), and the somatosensory map was adapted from (*55*). **(C**)Contours of the intrinsic imaging response peaks for individual deflection of the three whiskers Alpha, C2 and Delta overlayed on an image of surface blood vessels seen under green light through the cranial window. **(D)** Histology reconstruction of the barrel map overlayed on the same image of the cortical surface as in panel 1C. Note the overlap between functional and anatomical localizations of the barrels. **(E)** Left: Licks when the bar was in the green Rewarded zone were rewarded with water. Licks when the bar was in the red No lick zone triggered the end of the trial and the start of a 5 s intertrial interval. Right: Snapshots of the bar during a single trial. The trial always starts with the bar in the most caudal position. The bar moves according to a preloaded trajectory that enters at least once the Rewarded zone, and may reverse direction during the trial. Each trial ended either when the mouse licked in the No lick zone, or at the end of the trajectory (10-20 s). The color code for Rewarded and No lick zones is identical throughout all Figures. **(F)** Six example trajectories of the optogenetic stimulation. **(G)** Histogram of the times of first entry of the optogenetic stimulation in the Rewarded zone, for the 252 trajectories in the database. **(H)** Histogram of the average angular speed of the optogenetic stimulation, for the 252 trajectories in the database. The bar can move either clockwise or counterclockwise.

Mice were trained for 5 to 10 daily sessions of 30 minutes each. In Fig. 2A, the first 20 trials are shown for the first and last (10^th^) session for one mouse. During the first session, the mouse licked randomly and consequently was often punished with the immediate end of the trial. During the last session, the mouse successfully refrained from licking in the No lick zone. Instead, the mouse waited until approaching the Rewarded zone and then licked as it entered and crossed the Reward zone (Fig. 2A).

**Fig. 2.**
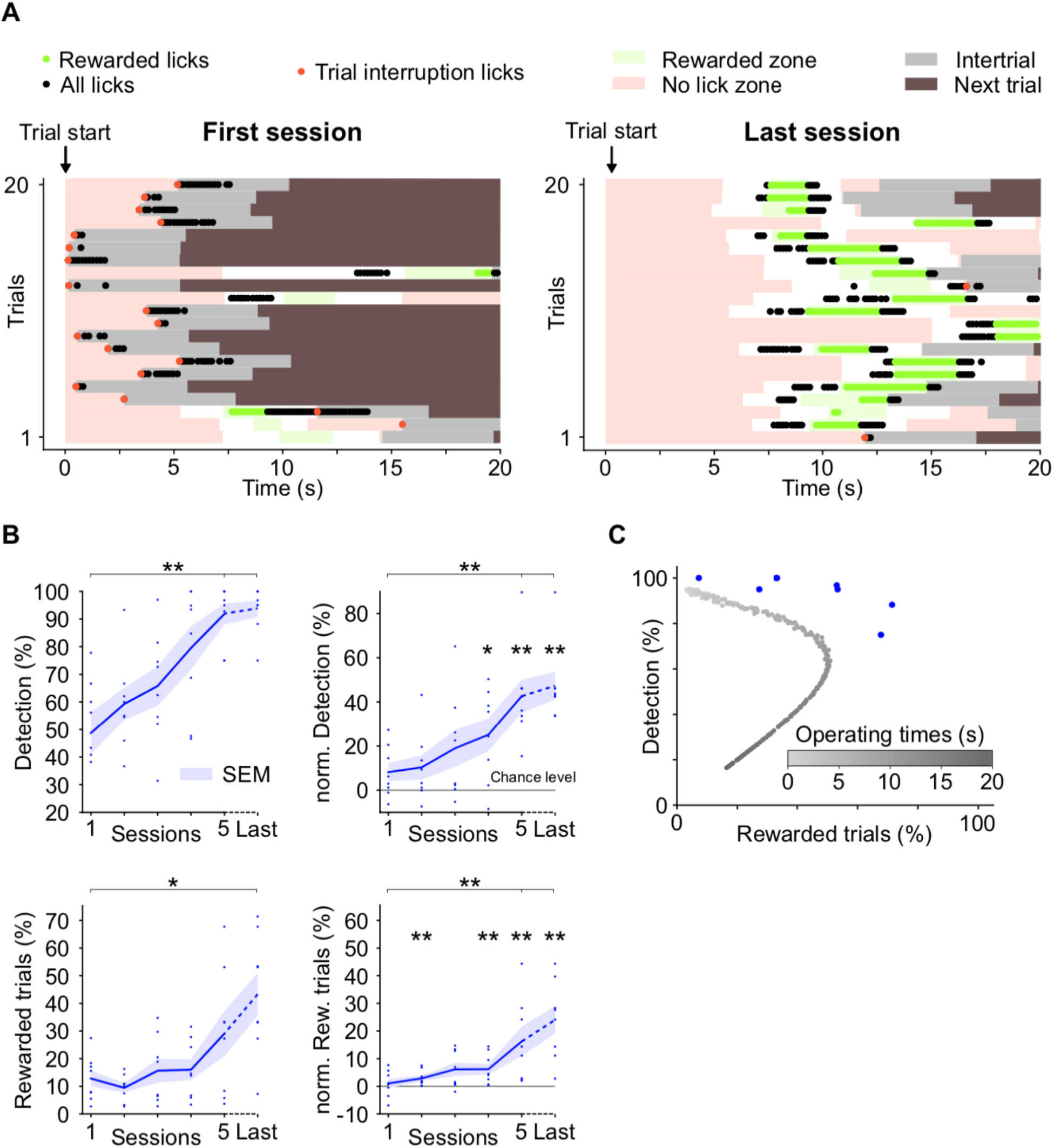
Mice learn to discriminate angular positions of a continuously rotating optogenetic stimulus. **(A)** Raster plot of licks (dots) during 20 consecutive trials in the first and tenth session for one mouse. Shading indicates either the zone in which the optogenetic bar is located during the trial (same color code as Figure 1), the inter-trial interval (grey) or the next trial (brown). Red dots indicate licks that occurred when the bar was in the No lick zone. Green dots indicate rewarded licks. **(B)** Average learning curves (+/− SEM, n = 8 mice) quantified by the Detection percentage (Top) and the percentage of Rewarded trials (Bottom). Left, raw performance measures. Right, chance level was calculated for each session by bootstrapping the trials (see Methods) and was subtracted from the raw performance measures to yield the normalized measures. Three mice were trained for more than 5 sessions (7, 10, 10). In each graph, the performance of the mice during their last session is labelled Last. Statistical tests between different sessions are Wilcoxon tests, while tests comparing performance to chance are one sample Wilcoxon tests. **(C)** Performance curve of an algorithm solving the task with a pure temporal strategy (see Methods). Each gray dot corresponds to the performance of the algorithm when it triggers licks starting at a specific time, called the operating time (grayscale). The blue dots indicate the mice performance during their last session. Statistical test: Wilcoxon test on the signed distance to the grey curve, p = 0.0078.

All 8 trained mice learned the task in 5 days (Fig. 2B). We quantified their performance by two metrics: the percentage of Rewarded trials, and the Detection level, which is the percentage of rewarded trials among the trials with available rewards. The average Detection level for the 8 mice increased across sessions, reaching over a 90 % score for the 5^th^ session (Fig. 2B Top Left, Wilcoxon signed-rank test, p = 0.0078, minimum value for n = 8). To check that this effect was not only due to changes in the frequency of licking, we estimated a chance level for each animal and each session by bootstrapping, which preserves the sequence of licks in time (see Methods). After subtracting this chance estimate from the Detection percentage, there still remains a significant increase of the Detection level across the 5 sessions (Fig. 2B Top Right, Wilcoxon test, p = 0.0078). This suggests that all animals learned to locate which sector of the stimulated cortical area leads to availability of reward. The average percentage of Rewarded trials increased more slowly and reached performance levels statistically above chance after 5 sessions (Fig. 2B Bottom right, one-sample Wilcoxon test, p = 0.0078). Compared to the Detection percentage, this performance measure also takes into account the fact that animals learned to refrain from licking in the No lick zone, thus leading to a decrease in the number of non-rewarded trials. Three animals out of 8 were trained for more than 5 days, and population analysis on all Last sessions showed that the performance of the animals continued to increase. Indeed, when comparing Last sessions to Sessions 5, the percentage of Rewarded trials was higher for those three mice.

We were concerned by the possibility that the mice learned the average timing from the onset of the trial to the entrance in the Rewarded zone, without relying on the angular position of the photostimulation. To estimate the performance of a mouse that would use this strategy, we designed an algorithm which solves the task using only time clues. The results from this algorithm are shown in Fig. 2C for all possible times of onsets of licks, called operating times. Each point corresponds to the algorithm’s performance (percentages of Detection and Rewarded trials) when submitted to the 252 trials for a given operating time (see Methods). Not surprisingly, short operating times (light grey points) resulted in most licks falling in the No lick zone, which in turn resulted in very low numbers of Rewarded trials despite good Detection values. Long operating times (dark grey points) led to late lick onsets, which also mostly missed the rewarded zones. Middle values could lead to better performance (Fig. 2C). However, all mice demonstrated higher performance than any version of the temporal algorithm (Wilcoxon, p = 0.0078). This comparison confirms that mice did indeed use the spatial location of the stimulation and its relation to the different behavioral zones to guide licking, and not only a temporal strategy.

To investigate this spatial aspect further, we analyzed the distribution of the stimulation angles at lick times. Fig. 3A shows the radial distribution of the photostimulation bar at the time of all licks (Top row) before and after learning, for the same mouse and sessions as in Fig. 2A. The proportion of licks for the Rewarded zone increased with training (from 19 % in session 1 to 61 % in session 10), while the proportion of licks for the No lick zone decreased (from 26 % to 1 %). Interestingly, when we looked specifically at the lick onsets (first licks of a rapid lick sequence, see Methods; Fig. 3A Bottom row), we observed that after learning, they occurred most frequently for the zones flanking the Rewarded zone (Bottom graphs in Fig. 3A; see also Fig. 2A, Right). This suggests that the mouse specifically started licking after the bar left the No lick zone and before it entered the Rewarded zone. Population analysis confirmed this redistribution of licks and of lick onsets across cortical photostimulation zones (Fig. 3B, n = 8 mice). Thus, mice learned to shape their licking behavior according to spatial cortical cues in order to increase the obtention of reward. This is also shown in Fig. 3C, in which we focused the analysis around the entry in the Rewarded zone by averaging trials entering from the left with trials entering from the right (symmetrized with respect to the rostro-caudal axis). In the naive animals, lick onsets were randomly distributed (Left), whereas the expert animals started licking well before the entry in the Rewarded zone (Right), and then kept licking until receiving rewards.

**Fig. 3.**
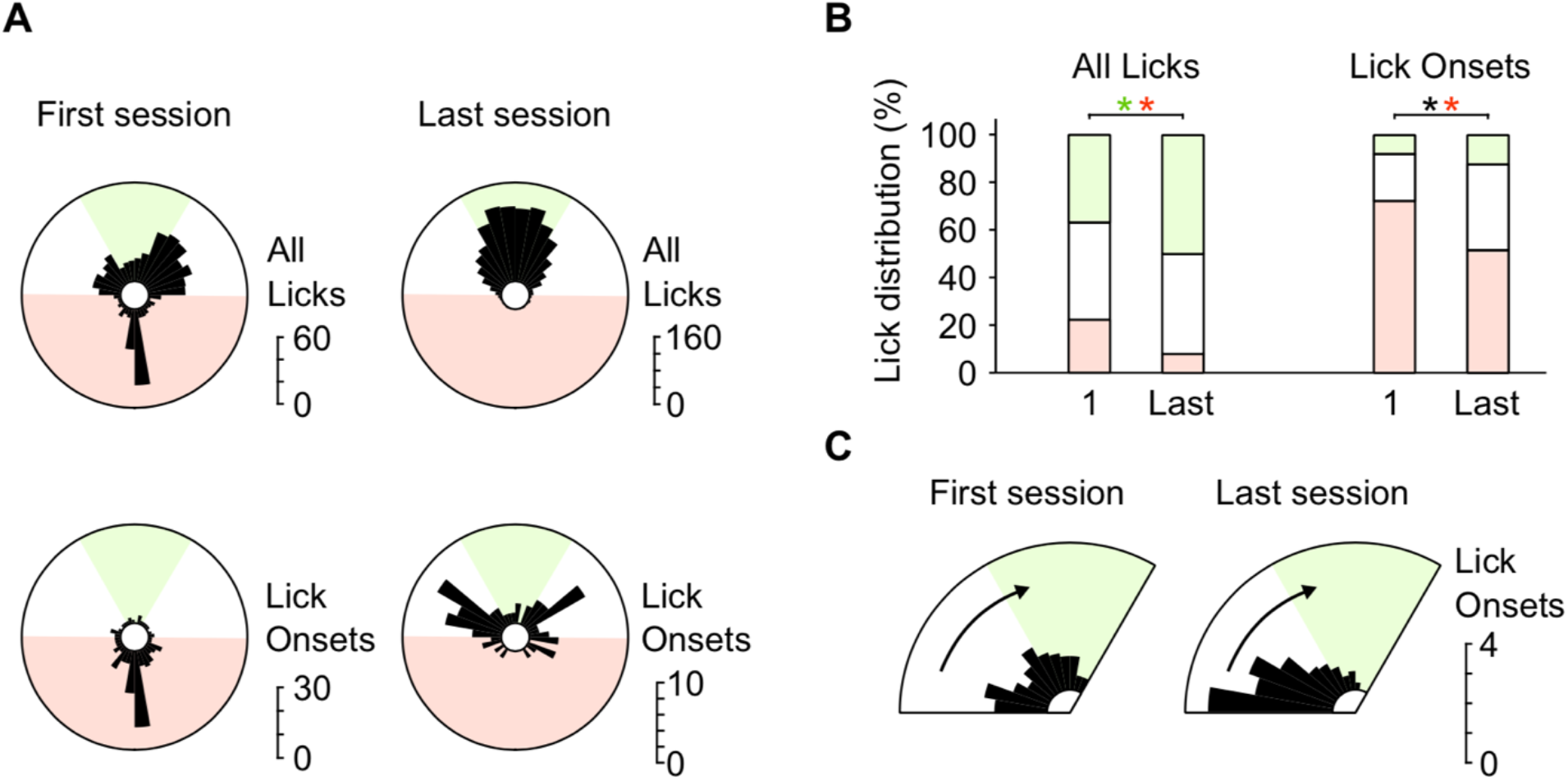
Redistribution of the angular positions of the optogenetic bar at lick times shows spatial anticipation of the reward. **(A)** Spatial distribution of the optogenetic stimulus angle for all licks (Top) and for lick onsets (Bottom, see Methods for lick onset definition), for the same mouse and sessions shown in Fig. 2A (First session on the Left, Last session on the Right). **(B)** Distribution of licks for which the optogenetic bar was in the different behavioral zones (green: Rewarded; red: No lick; white: neutral zones) averaged over all mice for their first and last session, for all licks and for lick onsets. Statistical tests: one-sided Wilcoxon tests, the black asterisk corresponds to the neutral zone. **(C)** Average spatial distribution of the stimulus angle for lick onsets around the entry in the Rewarded zone. Trials entering from the right were symmetrized so that all trials could be averaged together. Only the first 20 rewarded trials at most of each mouse for a given session were used for this analysis, ensuring that the mice were highly motivated.

To assess if mice were able to finely discriminate the different angles of stimulation, three mice which learned the task in 5 days were trained for 5 more days in a Difficult condition, in which the No lick zone was larger (Fig. 4A) to the detriment of the neutral zone. The first 20 trials of session 10 for a single mouse are shown in Fig. 4B. In 14 of these trials, the mouse refrained from licking until the bar left the No lick zone, and was then able to retrieve many rewards during the Rewarded zone traversal. On average, despite an initial drop in performance, these mice continued to increase their performance during the 5 more additional days of training in the Difficult condition (Fig. 4C). All three animals performed better than the temporal algorithm trained on this new task (Fig. 4D), indicating that again, they did not use a time-based strategy to solve the task. We computed the angular distribution of optogenetic patterns at lick onset times (Fig. 4E-G). When the Difficult condition was first introduced, the mice often started licking when the bar was in the No lick zone, particularly the zone where licking was allowed in the initial Standard protocol (Fig. 4F, orange area). Mice learned to avoid that zone, redistributing lick onsets so that by the last sessions, most occurred in the Rewarded zone or just before (Fig. 4E on an individual mouse, 4F-G average over 3 mice). Overall, these results show that mice, by relying on the continuity of the map in the barrel cortex, are able to discriminate finely different angles of the patterned optogenetic stimulation, even anticipating the sequence of movement from one zone to another.

**Fig. 4.**
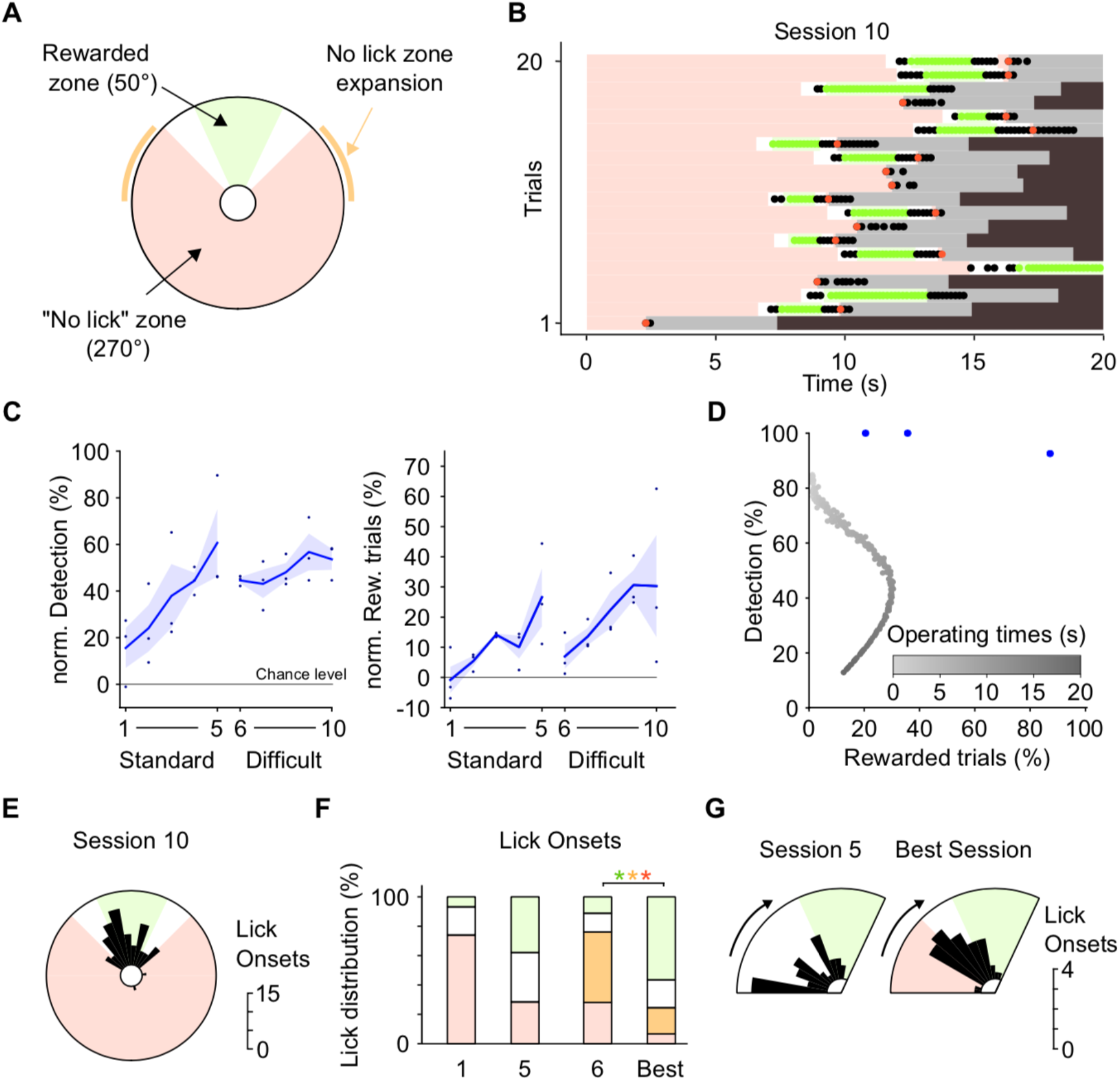
Mice can finely discriminate spatial zones of stimulation. **(A)** Rewarded and No lick zones for the Difficult condition. Orange lines: expanded area of the No lick zone. **(B)** Raster plot of licks (black and colored dots) during the first 20 trials of the last session of a mouse trained on the Difficult task for 5 days. Conventions are the same as for Fig. 2A. **(C)** Average learning curves (+/− SEM, n = 3 mice), quantified by the normalized percentage of Detection (Left), and the normalized percentage of Rewarded trials (Right). **(D)** Performance curve of an algorithm solving the Difficult condition task with a pure temporal strategy (see Methods and Fig. 2C). The blue dots indicate the mice raw performance during their last session. **(E)** Spatial distribution of the optogenetic stimulus angle for lick onsets for the mouse and session shown in panel B. **(F)** Distribution of lick onsets for which the optogenetic bar was in the different behavioral zones, averaged over all mice for the first and last session of the two training conditions. Green: Rewarded zone; Red: No lick zone of the Standard condition; Orange: No lick expansion zone for the Difficult condition; White: neutral zone. Best session corresponds to each mouse highest performance between session 9 and session 10. This was necessary because the task was very challenging, so performance was highly variable. Statistical test: one-sided MW test. **(G)** Average spatial distribution of the stimulus angle for lick onsets for trials during which the optostimulation entered the upper half of the disk, symmetrized for right entries as for Fig. 3C. For each mouse, only the first 20 of such events per session are represented.

### Mice use the spatial continuity of the stimulation space

To check if the anticipation of lick onsets shown in Fig. 3C and 4G is based on a spatial anticipation and not a pure temporal one, we trained 3 naive animals during 10 sessions each in a Shuffled condition (see Methods). The trajectories were still drawn randomly from the original database, but four angular sectors of the stimulation area were swapped (Fig. 5A). As a result, the optogenetic bar could now jump from one sector to a non-adjacent one when crossing one of the sector boundaries, breaking the spatiotemporal continuity of trajectories. The initial position and the Rewarded zone were unchanged. This task proved to be more challenging to the animals. Across the 10 sessions of training, the percentage of rewarded trials did not increase significantly, remaining far below values reached in the Standard condition (Fig. 5B, Right). Nonetheless, two out of three animals reached a 100 % Detection level after 10 days (Fig. 5B, Left). These results suggest that the animals had difficulty completing trials successfully because of the spatial discontinuities, but could nonetheless learn which area corresponded to the Rewarded zone as efficiently as mice in the Standard condition. Interestingly, when we looked at the time course of successful trials, we found that mice learning the task in the Shuffled condition developed a different strategy than in the Standard condition (Fig. 5C-E). Indeed, as seen on 20 completed trials (meaning not interrupted by licks) for one mouse (Fig. 5C), the animal started licking only when the optogenetic bar reached the Rewarded zone instead of anticipating its entry as in Fig. 2A. We quantified this observation by measuring the delay between the lick onsets and the optogenetic bar entry in the Rewarded zone (example in Fig. 5C), for sessions with a high Detection value (> 90 %, sessions E and more; one mouse was thus excluded from this analysis since it never reached 90 % in Detection). In the Standard condition, mice always anticipated the Rewarded zone entry by a median of ~1 second. In contrast, in the Shuffled condition they waited until after entry to start licking (Fig. 5D). This difference disappeared after 3 additional days of training, indicating that mice could eventually learn anticipation, despite the discontinuities in space. The initial lack of anticipation was confirmed by comparing the spatial distributions of the angular positions of the bar during lick onsets for mice which learned either the Standard (n = 8) or Shuffled (n = 2) condition (Fig. 5E, session E+1).

**Fig. 5.**
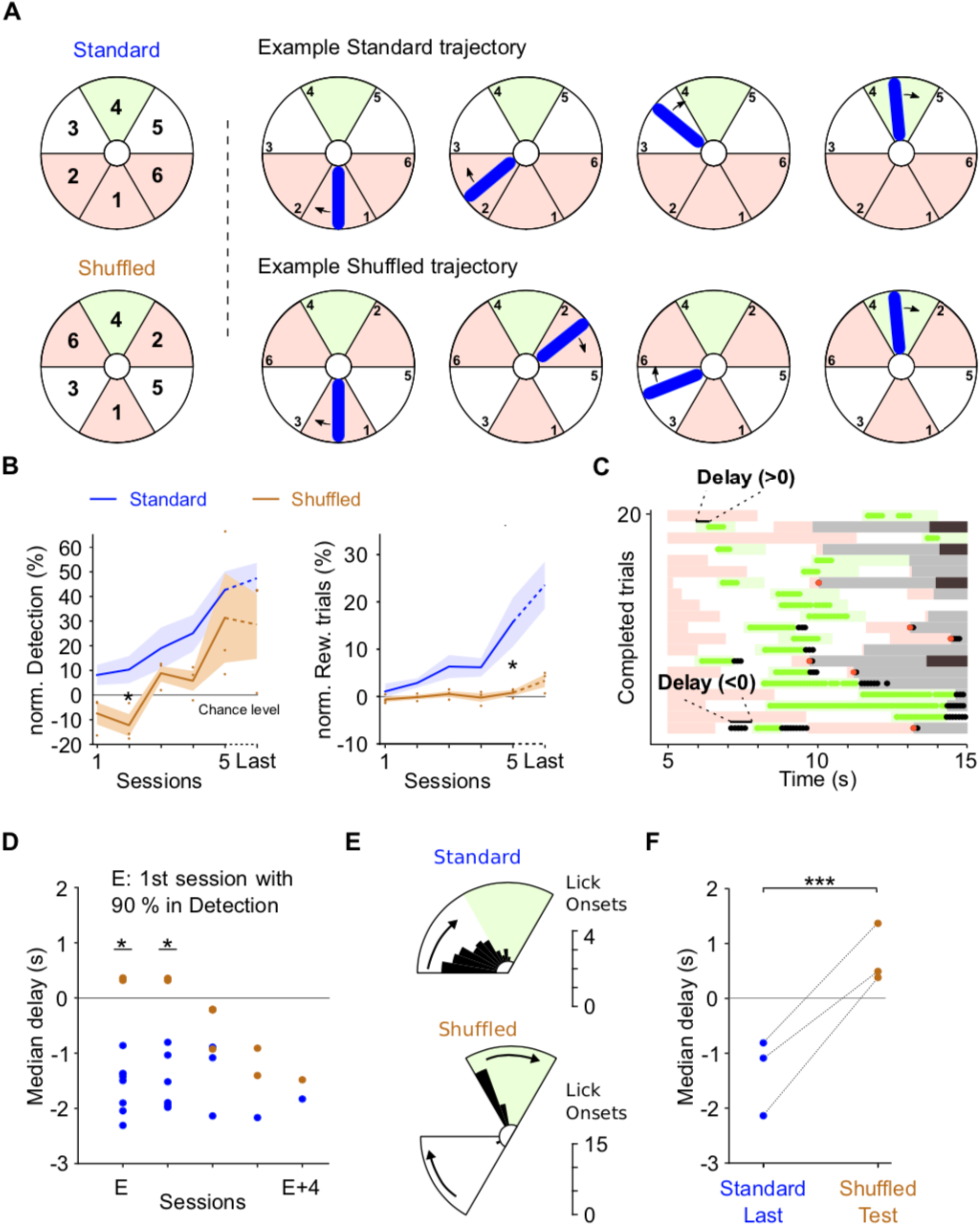
Mice rely on the spatial continuity of the stimulated area to learn the task. **(A)** Left, Reorganization of the angular sectors of the stimulation disk from the Standard condition (Top) to the Shuffled condition (Bottom). Sectors 1 and 4 are unchanged, other sectors are permuted as shown. Right, Example trajectory in the Standard condition (Top) and the same trajectory in the Shuffled condition (Bottom). **(B)** Average learning curves for the Standard (blue, n = 8 mice, 5-10 days; same as Fig. 2B) and Shuffled (brown, n = 3 mice, 10 days) conditions. Last session performance is labelled Last. Statistical test: two-sided MW between the two conditions, for each session. **(C)** Raster plot of 20 consecutive completed trials for a mouse trained in the Shuffled condition, for the first session with a high Detection percentage (session 6). The delay is the time from the entry in the Rewarded zone to the first reward. Only trials which were not interrupted by licks in the No lick zone are displayed. **(D)** Median delay from Reward zone entry to first rewarded lick as defined in panel C for the Standard (blue) and Shuffled (brown) conditions. For each mouse, Session E is the first session with a Detection rate of 90 % or more. Only consecutive sessions with a Detection rate of 90 % or more are shown. Statistical test: one-sided MW test on the delay medians, between the two groups of mice. **(E)** Average spatial distribution of the stimulus angle for lick onsets for rewarded trials, symmetrized for right entries as for Figure 3C, for both the Standard (n = 8) and Shuffled conditions (n = 2). We averaged the distributions of the second session with a detection rate over 90 % (E+1), across animals. One animal out of three in the Shuffled condition never reached a 90 % Detection level, thus was excluded from this analysis. Only the first 20 rewarded trials (at most) of the session are represented. **(F)** Median delay from Reward zone entry to first rewarded lick, for the last session of three animals trained in the Standard condition, and for a test session in the Shuffled condition one day later. Statistical test: one-sided MW on the distributions of delays between the two conditions for each mouse. Each of the three p-value is under 0.001.

We wondered whether animals that had already learned the task in the Standard condition could easily adapt to the Shuffled condition. Three expert animals in the Standard condition were tested in the Shuffled condition for 1 session. Despite a drop in performance, these animals remained experts at detecting the Rewarded zone (Detection = 100 %) but stopped anticipating the reward (Fig. 5F), demonstrating a direct effect of the spatial continuity of trajectories on behavior. To summarize, these results support the view that in learning to track the cortical optogenetic bar, the mice take advantage of the spatial continuity of the zones traversed on the barrel cortex to predict the future trajectory, and to start licking for reward shortly before its availability.

### Learning is disrupted by discontinuities in the cortical map

The continuous topographic organization of the barrel cortex might be critical for learning. Here, we trained naive mice to learn the same task while centering the photostimulation outside of the barrel cortex. Specifically, we first selected a cortical area where the representations of the trunk, the limb and the snout are adjacent (body S1, bS1), which results in discontinuities in the body map (Fig. 6A). This specific cortical area was localized using intrinsic imaging of the whisker barrels and of the forepaw (Fig. 6B). None of the three animals reached performance comparable to the animals that were trained with photostimulation on vS1, neither quantified by the percentage of Rewarded trials (two-sided Mann-Whitney (MW) test on the last sessions, p = 0.032) nor by the percentage of Detection (two-sided MW test, p = 0.019). Mice did not seem to distinguish the different stimulation zones, and produced seemingly random licking sequences. Second, to further explore this issue, we centered the stimulation on the posterior parietal cortex (PPC, 1.5 mm caudal to the C2 representation in vS1, Fig. 6A) in four animals already trained on S1. The rationale was to test learning of the photostimulation task for a non-primary sensory area, for which topography is not a strong organizing principle. Similar to the mice trained on bS1, these mice did not learn the task (Fig. 6D) neither. The significant difference in Detection between vS1 and PPC on the first sessions may be partially explained by the fast detection learning of vS1 mice already within the first session, and by the pre-training on a vS1 task for two PPC animals. Together, these results show that the continuous topography of the primary somatosensory cortex is necessary for learning the optogenetically mediated task.

**Fig. 6.**
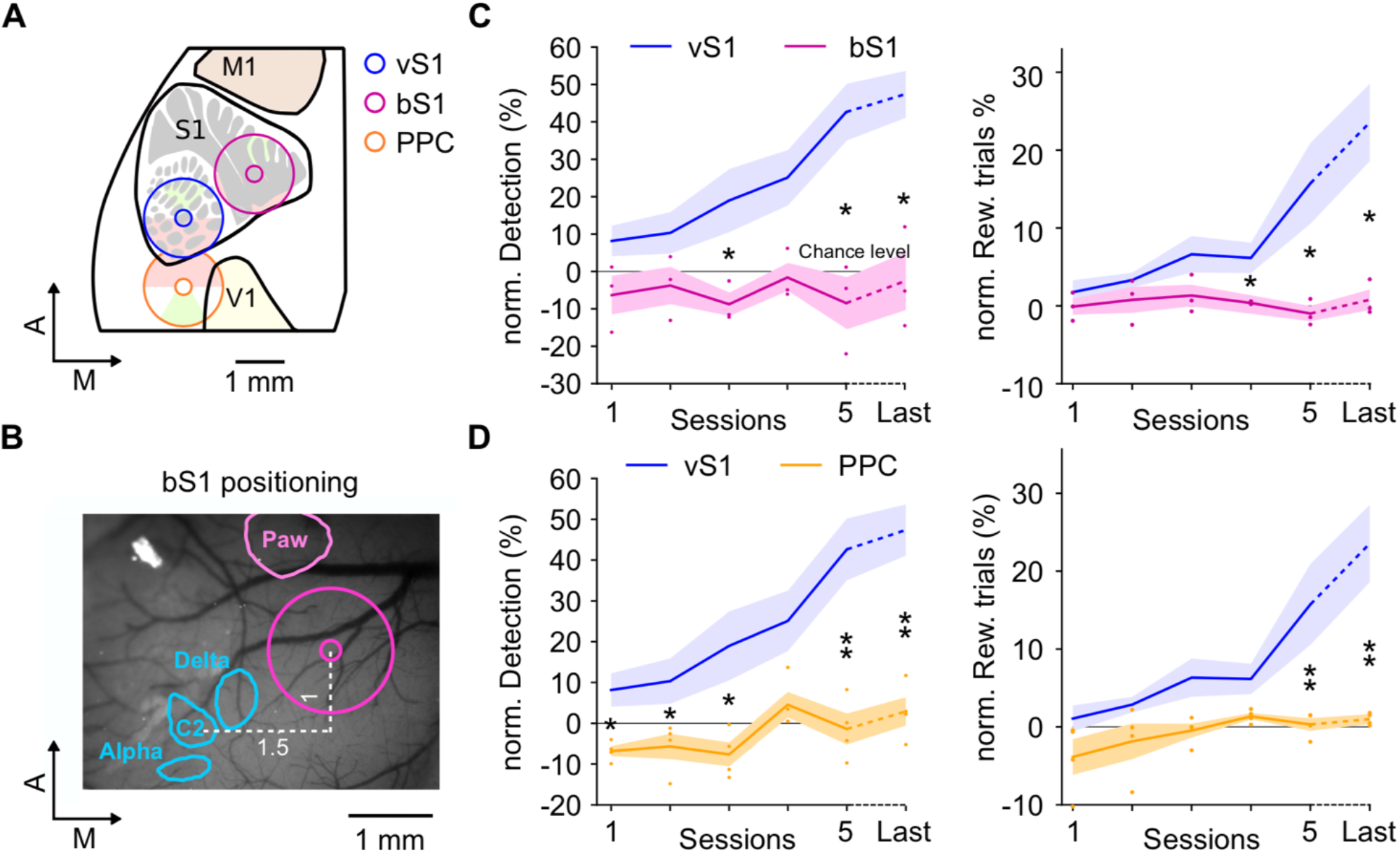
Learning is only possible in a cortical area that contains a continuous topographic representation of peripheral surface. **(A)** Location of the stimulation disk in in vS1 (blue), bS1 (magenta) and PPC (orange). The Rewarded and No lick zones are shown inside the disks. **(B)** Contours of the intrinsic imaging response peaks for individual deflection of the three whiskers Alpha, C2 and Delta (blue), and for mechanical vibration on the paw (pink), overlayed on an image of surface blood vessels seen under green light through the cranial window. The magenta circle corresponds to the location of the stimulation disk on bS1. **(C)** Average learning curves for stimulation of vS1 (blue, n = 8 mice, 5-10 days, same as Fig. 2B) and bS1 (magenta, n = 3 different mice, 10 days). Statistical test: two-sided MW between the two groups of mice, for each session. **(D)** Average learning curves for stimulation on vS1 (blue, n = 8 mice, 5-10 days, same as Fig. 2B), and on PPC (orange, n = 4 mice, 5-10 days). Two mice were trained during 10 days on the PPC and showed no learning. Two other animals were already expert after training on vS1 condition for 5 days, and were trained on the PPC for only 5 days during which their performance stayed at chance level. Same statistical test as in panel C.

## Discussion

Using a cortical stimulation protocol that induces activity sweeping on the cortical surface, we have shown that mice could learn to use patterned cortical stimulations as dynamical cues to obtain rewards. Learning strongly depended on the interplay between the spatiotemporal structure of the stimulus and the topography of the targeted cortical area. Overall, integration of the cortical inputs could take place within the time frame of our experiments only when the stimulus was continuous across space, and when the target area included a continuous topography of the sensory periphery.

Testing the functional significance of cortical topography in a perceptual task called for a high-resolution and flexible way to manipulate the neuronal activity across the cortical surface, a technical goal that is not easily achieved by electrical stimulation. The use of transgenic mice expressing channelrhodopsin in excitatory neurons of the cortex (*35*) allowed us to stimulate neurons by projecting a light pattern varying smoothly in space and time. Our particular choice of a rotating bar was guided by our long-term interest in encoding the angle of an articulation (*36*). In this context, the Rewarded zone corresponds to a target range of angles of the articulation. One important feature of optogenetic stimulation is that it activates predominantly neurons in the upper layers of the cortex (*37*). In vS1, as in visual cortex, the dense horizontal intracortical connectivity in these layers is a substrate for continuous propagating waves of depolarization (*38*, *39*). The fact that the mice were able to detect these supragranular mesoscopic patterns suggests that cortical patterns can indeed acquire functional relevance for perception and behavior. Interestingly, the delay between optogenetic stimulation and lick response was comparable to reaction times following peripheral stimulation. This could be observed for example when the mouse was not able to anticipate the patterns, as in the Shuffled condition, in which the mouse licked after the Rewarded zone entry (Figure 5). Reaction times started at around 300 ms, confirming earlier observations (*11*, *40*), and within the range of lick response times after a single whisker deflection (*19*, *41*).

During learning, mice first improved their task performance by licking reliably when the optogenetic bar entered in the Rewarded zone, thus increasing their Detection percentage up to 90-100 % already after five days. In parallel, the average percentage of Rewarded trials slowly increased but remained under 50 %, a direct consequence of the fact that most trials were still aborted because of licks in the No lick zone (Figure 3). This asymmetric dynamics between rewarded and non-rewarded trials is similar to the dynamics described in a Go/No-go discrimination task, in which the animal typically learns to lick for the S+ faster than it learns to refrain from licking for the S- (*42*).

Trajectories varied greatly in speed and direction (Figure 1), so that predicting the time of entry in the Rewarded zone could hardly be a successful strategy. We verified it by comparing the performance of the mice to that of time-counting algorithms (Figures 2 and 3). Each of the 8 mice was more successful than these algorithms, arguing against a pure temporal strategy. Moreover, mice could learn the task in the Difficult condition in which the temporal window for reward opportunity is shifting by several seconds from one trial to the next. It is also unlikely that the task was solved by a purely spatial strategy. Most often, mice anticipated the reward zone entry by licking shortly before, and in fact this turned out to be their preferred behavior. They adapted this anticipation to the condition they were tested in, with anticipatory licks less numerous and more finely timed when the No lick zone was expanded in the Difficult condition (Figure 4). When we suddenly introduced the Shuffled condition, expert animals were clearly perturbed by the discontinuous trajectory of the bar at the Rewarded zone entry, and first stopped anticipatory licking, preferring to trust only the ongoing spatial pattern of stimulation (Figure 5F). Furthermore, when naïve animals were trained repetitively in the Shuffled condition, learning was strongly impaired, as observed in the percentage of Rewarded trials which stayed very low (Figure 5B). However, these mice eventually learnt to anticipate the spatial jump and the upcoming reward opportunities, somehow adapting to the particular set of trajectories of the bar (Figure 5D). Overall, these results indicate that the animals use the spatiotemporal characteristics of the optogenetic bar to guide licking, particularly the continuous movement from one zone to the next.

The Shuffled condition induces a jump of activity from one cortical zone to another that is not spatially contiguous. This is similar to what happens following the surgical rotation of a skin flap, without severing nerve fibers (*43*). In those experiments, cortical topography was remodeled over the course of two weeks, so that a new somatosensory map underlined by modified receptive fields emerged, matching perfectly the new contiguity of peripheral skin zones. In fact, both maps observed before surgery and after experience-dependent remodeling co-existed at the end of the study. In our case, non-contiguous cortical patterns cannot remap because we manipulate them directly and not via peripheral afferents. Nonetheless, we hypothesize that a reorganization of spatial readout mechanisms by upstream areas could be at play. This would explain that mice trained in the Shuffled condition could learn to anticipate reward opportunities, despite overall poor performance.

Disrupting the continuity of the perceptual experience could be achieved either by breaking the continuity of the cortical stimulus as discussed above, or by applying stimulation on an area in which topography is not continuous.

When we tested continuous light patterns applied on the discontinuous topography of body S1, mice could not learn the task in 10 days of training, not even showing improvement in detection of the Rewarded zone (Figure 6). This was quite surprising, and we do not exclude that learning could be possible with more training time. We interpret their lack of performance as related to the fact that the optogenetic bar traverses cortical zones that correspond to non-contiguous peripheral skin zones. Thus, the stimulation could be perceived as a stimulus jumping between many different locations on the contralateral limbs and trunk, including a fragmented perception when the bar simultaneously excites distinct body zones. If the somatosensory cortex is used by the animal as a body model to simulate ongoing behavior and its consequences (*30*), our induced patterns of activity on bS1 could indeed be difficult to interpret by the animals. The same argument could explain the lack of learning when stimulation targeted the PPC. Overall, our findings thus fit with the general idea that the topography of a cortical area shapes the integration of activity by neuronal networks, whether induced directly at the cortical level or resulting from peripheral stimulation.

Understanding how an animal may extract information from ongoing cortical patterns is an active area of research, with immediate relevance to the field of neuroprostheses. Recent studies question whether cortical stimulation should aim at biomimicry, by making the spatial and temporal aspects of the artificial stimulation as close as possible to the cortical response inferred by the targeted natural stimulus (*44*, *45*). In the somatosensory system, the main objective has been to restore the sense of touch or proprioception, either to reach a fine dexterity in a closed loop context or to elicit sensory percepts and alleviate sensory deficits. In the visual system, the importance of biomimicry has also been investigated (*46*). However, most previous studies relied on discrete patterns of stimulation in space and time, whereas everyday prosthesis use will generate complex and continuous sensory feedback. In this work, we designed a novel stimulation continuous in time and space across the cortical surface. We do not argue that this is a biomimetic activation, but it shares some characteristics with known activity patterns in the cortex, notably the propagation of cortical waves of activation. Our results indicate that stimulation of a topographically-organized sensory cortex can be integrated with a high degree of spatial precision into behavior, including anticipatory processes which could promote dexterous movements of a closed-loop neuroprosthesis.

On longer time scales, it has been argued that plasticity mechanisms reorganize sensory cortex in order to match the structure of peripheral stimulation (*43*, *47*), a phenomenon that could also exist following neuroprosthesis use. However, the extent of such reorganization is still debated (*45*). Recent studies on participants using a sensorimotor bionic arm suggest that the location of the sensor on the bionic hand, and the tactile experience following electrical nerve stimulation, should roughly match at the time of nerve connection for long-term perceptual alignment of prosthesis and missing limb (*48*–*50*). Thus, remapping in adults would occur within certain limits (*44*), as has been described in details in the barn owl auditory localization system (*51*). A similar consensus has emerged in motor control of neuroprostheses, arguing that imposing a motor decoder from scratch takes much longer to learn than refining pre-existing cortical representations (*52*, *53*).

In conclusion, we showed that dynamical patterned cortical stimulation could be used by mice to drive licking for reward. Successful processing of the stimulus required that its spatio-temporal characteristics remained geometrically coherent with the known topographical cortical map. Notably, discontinuities in the map or in the stimulus impaired learning. Our results can help guide the design of sensory rehabilitation devices, allowing to track continuously the state of natural or artificial limbs.

## Materials and methods

### Mice

We used 6-week-old Ai-27 x EMX-Cre mice, expressing channelrhodopsin in excitatory neurons across the cortex (*35*). Experimental procedures have been approved by the French Ministry of Research and Ethics Committee #59 as part of project #858-2015060516116339. A total of 14 mice were successfully implanted, water restricted, and then trained in the task. During the training period, the mice only had access to water during the sessions as reward, and right after the session for supplementation whenever necessary.

### Mouse preparation

Surgeries were performed on anesthetized mice (1-4 % isoflurane anesthesia in 100 % air) placed on a regulated heating pad. The state of the anesthesia was assessed by breathing rate and response to tail pinch. The scalp was resected after lidocaine analgesia (200 mg/L, 0.1 mL) and conjunctive tissues were removed. A head-post was glued (cyanoacrylate glue) to the skull, then reinforced with dental cement. A 5 to 6 mm diameter craniotomy was then performed while preserving the dura, centered either on the stereotaxic coordinates of the C2 barrel in the primary somatosensory cortex (P1.5-L3.3 mm), or on a more medial location in between the paw representation and the barrel cortex (P0.5-L2.3 mm). A glass optical window of diameter 5 or 6 mm was glued to the borders of the craniotomy. The remaining exposed skull was covered with dental cement. At the end of the surgery, we administrated subcutaneously an analgesic (2 mg/mL Metacam, 0.1 mL) and a diluted antibiotic (2.4 % Borgal, 0.2 mL). Intrinsic imaging sessions through the window were performed 5 to 10 days after the surgery. During an imaging session, either a single whisker or the right forepaw was stimulated with a piezoelectric bender (Physics Instruments) 100 Hz, 5 ms square wave deflection) while red light (625 nm) was projected on the window right below light saturation. A CCD camera acquired 659*494 px images at a rate of 60 fps. The images were analyzed for space-time fluctuations in luminescence (Optimage, Thomas Deneux, NeuroPSI). These intrinsic imaging sessions were used to locate the C2, Alpha and Delta barrel locations, as well as the forepaw location (Fig. 1C & 6B). More details can be found in (*40*).

### Optogenetic photostimulation

During training, optogenetic stimulation was performed through the optical window (Fig. 1A) using a Digital Light Processing module (DLP, Vialux V-7001, 462 nm blue LED. Stimulation patterns consisted of light bars 700 microns long and 150 microns wide, rotating on a disk of diameter 1.5 mm. The disk was centered on the C2 barrel location (vS1, Fig. 1B, n = 11 mice). In three additional mice, the stimulation was centered on a point 1.5 mm medial and 1 mm rostral to the C2 barrel, thus on the trunk and legs representation (“body” S1, bS1, Fig. 6A-B). Among these 14 mice, four were subsequently trained with the stimulation centered on the posterior parietal cortex (PPC), 1.5 mm caudal to the C2 barrel (Fig. 6A; three mice after vS1 training, one after bS1 training). To avoid overstimulation of cortical areas, the center of the stimulated disk was never illuminated (white spot in Fig. 1D). Photostimulation was done at high power (measured, 10-15 mW per mm^2^). Spiking activity resulting from such photostimulation was demonstrated previously (*10*, *40*). Using a photodetector, we ensured that the edge of the photostimulation was sharp: intensity 5 % at 20 microns.

### Behavioral training

After surgery and intrinsic imaging, the mice were water restricted. Training started two days later. During the first session, the mouse was habituated to head-fixation and learned to lick water from a small tube. During this session, licks always triggered water rewards, and nothing happened if the mouse did not lick. Each subsequent session lasted 30 minutes, during which a randomized set of trials was presented to the mouse. Trials were separated by 5 seconds.

In the Standard condition, each trial consisted in the presentation of one trajectory of the photostimulation bar, starting from the most caudal position, and then rotating towards a rostral Rewarded zone with different dynamics (Fig. 1E-H). The bar angular position was updated every 10 ms. Each trajectory was taken from a database of 252 pre-loaded trajectories. These were obtained from a closed-loop BMI study performed with different mice (*36*). In that study, the activity of motor cortex neurons was used to drive the rotating bar. From the initial full dataset, we kept only the 126 trajectories lasting between 10 and 20 seconds, and entering at least once the Rewarded zone (green zone, Fig. 1E). In order to remove a possible right/left bias, for each trajectory, its symmetric trajectory with respect to the rostro-caudal axis was added to the database, yielding the full set of 252 smooth trajectories with evolving dynamics (Fig. 1F-H).

A lick had different consequences depending on the angular location of the photostimulation bar at that time. A lick when the light bar was in the Rewarded zone (green area, Fig. 1E) led to an immediate 10 μL water reward. A lick in the No lick zone (red area) immediately ended the trial and was followed by the 5 s intertrial interval, during which the cortex was not photostimulated. A new trial started immediately after. A lick in the neutral zones (white areas) had no consequence. If the animal drank more than 3 mL of water during one session, only one lick out of two was rewarded with water during the following sessions, starting with the first lick inside the Rewarded zone. The Rewarded zone spanned 60°, except for three mice for which it spanned 50°.

Three animals initially trained for 5 days in the Standard condition on vS1 were then trained for 5 days in a Difficult condition, in which the size of the No lick zone was reduced so that the task became more challenging (Fig. 4A, these were the three mice trained with a 50° Rewarded zone). Three other animals were also initially trained in the Standard condition and were then tested for one session in a Shuffled condition (Fig. 5A-F), in which the contiguity of the cortical sectors crossed during the trajectories was modified. Specifically, the stimulation disk was divided in six 60° sectors, and these sectors were swapped so that the trajectory jumped from one sector to another at the boundaries. The starting position and Rewarded zone were unchanged. Another group of three naive mice were directly trained in the Shuffled condition on vS1 (Fig. 5B-E).

Mice were usually trained every day for 10 days once enrolled in a protocol. However, in the first experiments, training was often stopped after 5 days of vS1 stimulation in the Standard condition because the performance of the animals was already very high. When training duration varied across animals, the last session of each mouse was labelled Last for group analysis.

### Histology

After training, mice were deeply anesthetized with isoflurane (4-5 %), euthanized by cervical dislocation, and perfused with para-formaldehyde (PFA). The brains were stored in PBS for two days or more, and then S1 tangential slices (100 microns thick) were cut and stained with cytochrome C oxidase. The alignment of the stained barrels and the blood vessels were computed using a homemade software (*34*). Briefly, this software uses the position of the transversal vessels to realign each slice with respect to each other. At the end of the process, the slice showing the blood vessels on the surface of the cortex and the slices showing the stained barrels could be superimposed accurately (Fig. 1D).

### Statistical analysis

We quantified performance by looking at the percentage of trials that were rewarded in each session. We also computed a Detection value (Fig. 2B), corresponding to the percentage of entries into the rewarded zone for which the mouse obtained at least one reward. Therefore, the Detection was not affected by trials that were interrupted before reaching the rewarded area. Overall, a high Detection value indicates that the mouse learned to lick for the Rewarded zone, while a high percentage of Rewarded trials requires that in addition, the mouse learned to refrain from licking in the No lick zone. These two measures of performance were normalized session by session by estimating a chance level obtained by bootstrapping. For each session, one hundred shuffled sessions were generated by loading random sequences of trajectories from the database while keeping the temporal sequence of licks from the real session. The simulated protocol followed all the task rules: if during a simulated trajectory, a lick happened while photostimulation was in the No Lick zone, a 5 s Intertrial interval was enforced and a new trial was loaded thereafter. The average performance of these simulations was then subtracted from the performance of the real session to obtain the normalized performance (Fig. 2B, 4C, 5B, 6C-D).

To demonstrate that mice were using spatial information from the cortical stimulus and not only a temporal strategy, we designed an algorithm that solves the task exclusively by using time elapsed in the trial. In the simplest version, this algorithm waits a fixed amount of time, called its operating time, then licks continuously until the end of the trial. In the version we used, instead of a fixed value, we picked waiting times randomly from a Gaussian distribution centered on the operating time, and with a standard deviation equal to the smallest standard deviation of response times observed in a trained animal. We generated one such algorithm for each operating time from 0 to 20 s (with 0.01 s steps). For each operating time, we quantified the performance of the algorithm by the Detection value and percentage of rewarded trials for a full session comprised of the 252 different trials.

To quantify the behavioral licking response of the mice as a function of cortical location, we analyzed the distribution of angular positions of the optogenetic bar at lick times (Fig. 3A). However, as a lick in the Rewarded zone leads to a reward and thus to more licks, these plots often showed huge numbers of licks not necessarily linked to the simultaneous photostimulation location. To disambiguate first licks from others, we defined an onset lick as a lick that was not preceded by another lick in the last 3 seconds.

Each statistical test used is described in the text and/or in the figure legends. *: P < 0.05. **: P < 0.01. ***: < 0.001. Unless stated otherwise, all tests are two-sided.

## Acknowledgments

We thank Guillaume Hucher for the histology, Esther Fournel for optogenetic characterization, Isabelle Ferezou for advice throughout the project, and Aurélie Daret for help with animal experiments and lab managing.

## Funding

We thank ANR Neurowhisk ANR-14-CE24-0019, FRM DEQ20170336761, Lidex iCODE and NeuroSaclay in the IDEX Paris-Saclay ANR-11-IDEX-0003-02, FRC AAP2018 and CNRS80.

## Author contributions

DES, LE and VE initiated the project, HL, DG, DES, LE and VE designed the experiments, HL performed the experiments with help from DG, HL analyzed the data with support from VE and LE, HL wrote the paper with inputs from all authors.

## Competing interests

The authors declare no competing interests.

## Data and material availability

All data needed to evaluate the conclusions in the paper are present in the paper and/or the Supplementary Materials. Additional data related to this paper may be requested from the authors.

## Notes

### Competing Interest Statement

The authors have declared no competing interest.

